# The X chromosome is a potential polarising signal for asymmetric cell divisions in meiotic cells of a nematode

**DOI:** 10.1101/2022.03.15.484444

**Authors:** Talal Alyazeedi, Emily Xu, Jasmin Kaur, Diane Shakes, Andre Pires-daSilva

## Abstract

The unequal partition of molecules and organelles during cell division results in daughter cells with different fates. Asymmetric cell divisions have been best characterised in systems in which extrinsic signals polarise the mother cell during cell division. However, the mechanisms of asymmetric cell division mediated by intrinsic signals, and the nature of these signals, are mostly unknown. Here we report an asymmetric cell division in the nematode *Auanema rhodensis* that may be cued by the X chromosome. In the wildtype XO male, the spermatocyte divides asymmetrically to generate X-bearing spermatids that inherit components necessary for sperm viability, and nullo-spermatids that inherits components to be discarded. We found that in XX mutant pseudomales, sperm components co-segregate with the X chromosome, supporting the hypothesis that the X chromosome is employed as a polarising signal for partitioning essential cytoplasmic components for sperm function.

## INTRODUCTION

An asymmetric cell division (ACD) is a division that results in sister cells with different sizes, morphology, chemical composition, or gene expression. ACDs are present in both unicellular and multicellular organisms, are crucial for embryonic development, cellular differentiation, and production of cell diversity (Gönczy, 2008; Venkei and Yamashita, 2018), whereas their dysregulation may result in tumorigenesis (Knoblich, 2010; Neumuller and Knoblich, 2009).

The polarity of a dividing cell is established by extrinsic or intrinsic factors. The best-known examples are the cases in which extrinsic signals provided by neighbouring cells polarise the dividing cell (Fuller and Spradling, 2007). Relatively little is known about intrinsic mechanisms controlling the polarity of an asymmetric dividing cell (Freisinger et al., 2013). We have previously described a cell division that undergoes asymmetric segregation that is likely to be guided by internal mechanisms. This division occurs during the final division of the male spermatogenesis in the nematode *Auanema rhodensis* (aka, *Rhabditis* sp. SB347) (Shakes et al., 2011; Winter et al., 2017).

Spermatogenesis is very well characterized in the nematode *Caenorhabditis elegans*. During the maturation of the spermatids, non-essential cytoplasmic components are extruded into vesicles named residual bodies (Fig. 1A). Thus, the final product of an XO spermatocyte division and differentiation is the formation of four spermatids, two spermatids with no X chromosomes (nullo-X spermatids) and two spermatids with one X chromosome each (X-bearing spermatids) (Fig. 1A). Unexpectedly, spermatogenesis in *A. rhodensis* results in only two viable gametes, each containing either one X chromosome (XO male, Fig. 1B) or two X chromosomes (Shakes et al., 2011; Tandonnet et al., 2018) (XX hermaphrodite, Fig. 1C). In *A. rhodensis* the non-essential sperm components are discarded into the nullo-X spermatids because of asymmetric partitioning of the cytoplasm during anaphase II (Fig. 1D) (Shakes et al., 2011; Winter et al., 2017). Thus, the nullo-X spermatids become cellular residual bodies.

**Figure 1.**
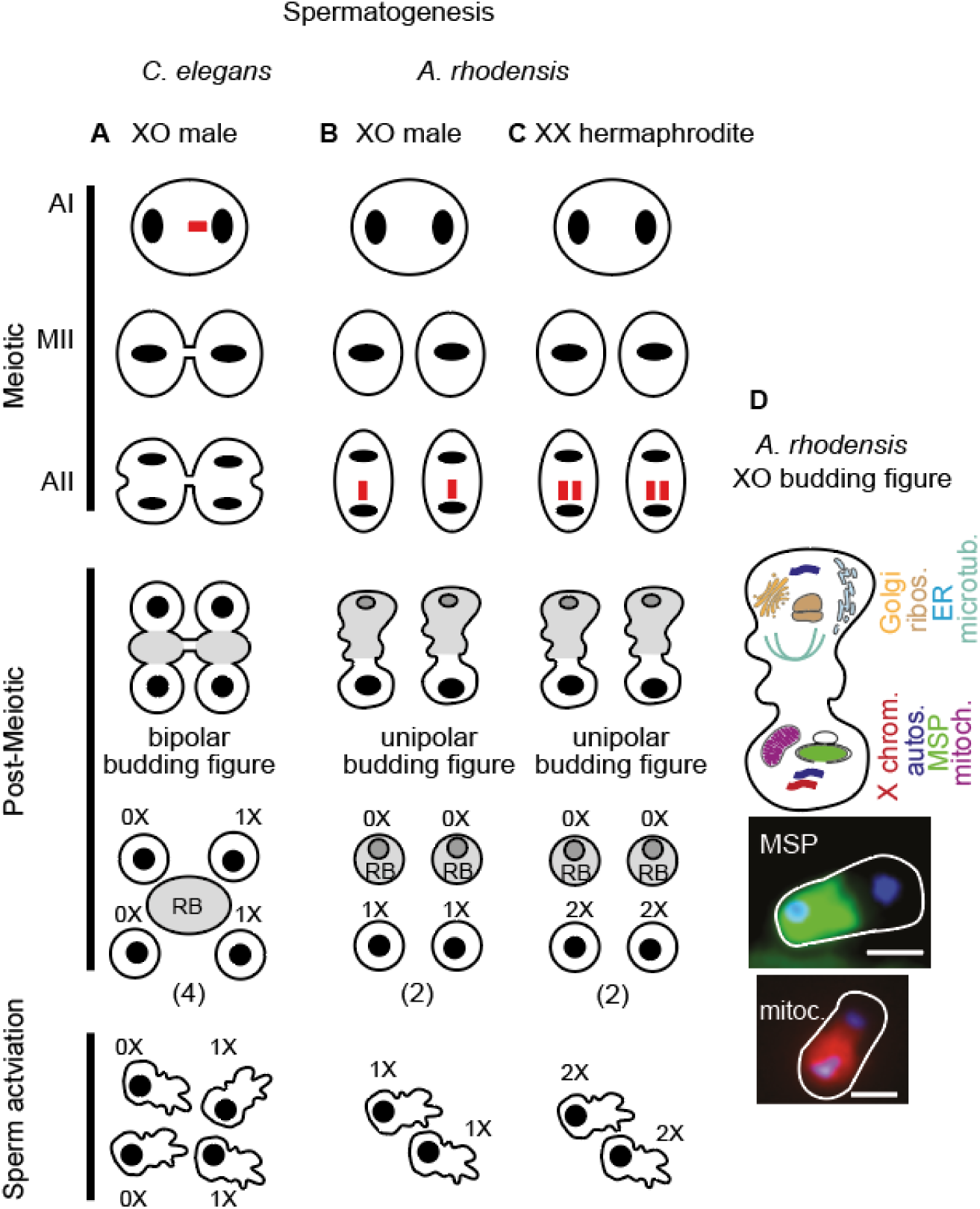
Comparison of spermatogenesis between *C. elegans* and *A. rhodensis*. In *C. elegans* males (**A**), the unpaired X chromosome (in red) lags during anaphase I, leading to the formation of two secondary spermatocytes that divide symmetrically to generate either two gametes with 1X or two gametes containing only autosomes (0X). A central residual body (RB) is formed in the postmeiotic phase. In *A. rhodensis* males (**B**), the sister chromatids of the X chromosome separate prematurely in meiosis I and the X chromosome lags during anaphase II. High rates of non-disjunction in *A. rhodensis* hermaphrodite spermatogenesis leads to the formation of 2X sperm (**C**). Only two functional gametes are formed during spermatogenesis of *A. rhodensis* (**B, C**), with the nullo-X sperm acting as a residual body. Cytoplasmic components partition during the second meiotic division (**D**), whereby specific organelles and proteins partition in opposite directions. Mitochondria and the sperm cytoskeleton protein Major Sperm Protein (MSP) partition with the X chromosome to one of the poles of the cell, whereas ribosomes, Golgi complex, and microtubules migrate to the opposite side of the dividing cell, which has only autosomes (Winter et al., 2017).

We hypothesized that the *A. rhodensis* X chromosome acts as a polarizing signal, guiding the partitioning of sperm-specific cytoplasmic contents to one of the poles of the dividing secondary spermatocyte. As a result of this ACD during spermatogenesis, *A. rhodensis* males and other species of the same genus produce mostly X-bearing sperm (Chaudhuri et al., 2015; Shakes et al., 2011; Winter et al., 2017). Thus, crosses between males and females result mostly in XX progeny, which can be either female or hermaphrodite (Chaudhuri et al., 2011a). Hermaphrodites, which produce nullo-X oocytes and XX-sperm, produce mostly XX self-progeny. To determine if the X chromosome acts as a polarizing signal, we isolated a sex determination mutant that is phenotypic male but that harbours two X chromosomes. We show that sperm components co-segregate with the X chromosome, further supporting that this chromosome acts as a polarizing signal.

## MATERIALS AND METHODS

### Nematodes strains and cultures

The *Auanema rhodensis* inbred strains APS4 and APS6 (Tandonnet et al., 2018) were maintained according to standard conditions for *C. elegans*, at 20 °C (Stiernagle, 2006). Nematodes were cultured on NGM plates seeded with the *Escherichia coli* strain OP50 or with the *E. coli* streptomycin-resistant strain OP50-1.

### Mutagenesis

*A. rhodensis* APS4 was mutagenized with the chemical mutagen ethyl methanesulfonate (EMS), as previously described (Chaudhuri et al., 2011b; Pires-daSilva and Sommer, 2004). To screen for a masculinizing phenotype (XX pseudomales), we isolated 521 F1 hermaphrodites derived from mutagenized P0s and transferred them to single plates to let them self-fertilize. To simplify screening for mutants that generate high rates of (pseudo)males, we screened for F2s derived from 3-day old hermaphrodites. We adopted this procedure because hermaphrodites of this age tend to generate less XO self-offspring (∼3%) than younger hermaphrodites (∼8%) (Chaudhuri et al., 2015). From plates in which potential pseudomale mutants were found (containing ∼25% of phenotypic males), 10-15 sister hermaphrodites were isolated to single plates to maintain the mutations as a heterozygous strain. Heterozygous hermaphrodites were selected based on the production of excess male progeny, which is consistent with the anticipated production of 25% XX male progeny

### *A. rhodensis* masculiniser (*Arh-mas-1*) crosses

The *A. rhodensis* masculiniser was named *Arh-mas-1*(*brz-3*), following the nomenclature described in Wormbase (www.wormbase.org). *Arh-mas-1*(*brz-3*) was backcrossed with the wildtype APS4 for three generations to remove background mutations generated during the mutagenesis.

To ascertain that the pseudo male has two X chromosomes, we crossed it to the polymorphic strain APS6. *Arh-mas-1* crosses with APS6 females were performed for ∼24 h at 20 °C. To isolate females for crosses, we left a 1-day old hermaphrodite to lay eggs, which were picked to individual wells of a 24-well plate. After 72 hours, those eggs develop into males, females or hermaphrodites. Females were distinguished from males by their tail morphology, and from hermaphrodites by their inability to self-fertilize (Kanzaki et al., 2017).

### Single nematode genotyping

The X chromosome genetic markers (markers 9686 and 12469) generated for *A. rhodensis* (Tandonnet et al., 2019), together with the primer sequences, restriction enzymes and fragment sizes are detailed in https://data.mendeley.com/datasets/63d7rrrx28/3#file-16ff094d-6c74-478a-a3f5-8878e89fd72f.

### Antibody staining

To detect the ER, we used an antibody against cytochrome P450 (CYPP33-E1). For mitochondria, we used an antibody against the beta-subunit of ATP synthase.

## RESULTS

### *Arh-mas-1* has a male phenotype and XX karyotype

We performed chemical mutagenesis in *A. rhodensis* to screen for a masculinizing sex determination mutant. This mutant would allow us to determine whether the X chromosome acts as a signal to partition the cytoplasm in dividing spermatocytes. If this hypothesis is correct, the expectation was that the two homologous X chromosomes segregate to opposite poles in the first and second meiotic divisions, following a Mendelian pattern (Fig. 2A, Model 1), to generate viable 1X spermatids. Although *A. rhodensis* XX females follow the canonical meiosis, XX hermaphrodites do not: the X chromosomes undergo premature chromatid separation in the first meiotic division, forming viable 2X spermatids and non-viable nullo-X spermatids (Tandonnet et al., 2018). Thus, similar to hermaphrodites, pseudomales may follow the pattern found in hermaphrodite spermatogenesis (Fig. 2A, Model 2).

**Figure 2.**
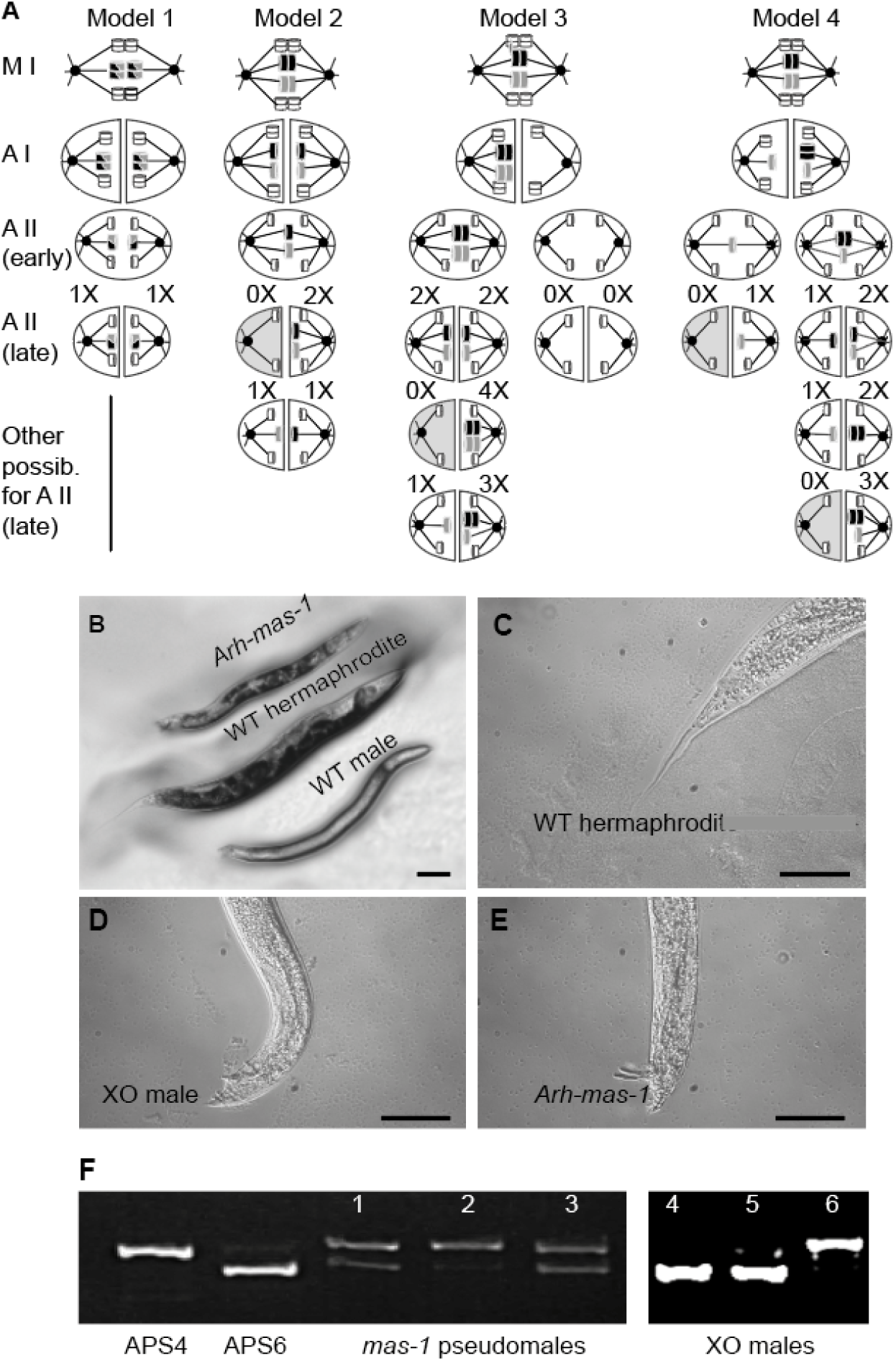
*Arh-mas-1* masculinizes XX animals. (**A)** Models for possible meiotic events in XX pseudomales. Homologous X chromosomes are in grey and black, and autosomes are in white. In the models, nullo-X sperm are discarded as residual bodies (grey cytoplasm) when the sister cell has an X chromosome(s). (**B)** The *Arh-mas-1* pseudomale has a gut dark pigmentation pattern distinct from the wildtype male. Bar, 10 μm. The tail of the hermaphrodite **(C)** is long and slender, whereas the male **(D)** and *Arh-mas-1* **(E)** are blunt and with spicules. Bar, 50 μm. **(F)** Genotyping of the X chromosome (chromosome marker 9686) in animals derived from crosses between pseudomales (APS4 background) and females (APS6).

Alternatively, those X chromatids may segregate equally to each pole to make viable 1X sperm. A third possibility is that the two X chromosomes segregate to one pole in the first meiotic division, generating spermatocytes that either make only nullo-X spermatids or 0-4X spermatids. (Fig. 2A, Model 3). The fourth possibility is a mixture of Models 2 and 3. The hypothesis of the X chromosome acting as a polarising signal for cytoplasmic partitioning would be falsified in case the cellular components important for post-meiotic sperm do not co-segregate with the X chromosomes in anaphase II.

We found an XX *A. rhodensis* sex determination mutant (*Arh-mas-1*) with a male morphological phenotype that is almost indistinguishable from XO wildtype males (Fig. 2B-E). Adult *Arh-mas-1* XX pseudomales that are three days or older display a gut pigmentation pattern distinct from XO males (Fig. 2B), which we used as a marker to distinguish between the two karyotypes. *Arh-mas-1* pseudomales do not show signs of partial feminization, as encountered in similar mutants in other nematode species (Hill et al., 2006; Hodgkin, 1979; Pires-daSilva and Sommer, 2004). *Arh-mas-1* pseudomales display mating behaviours and produce viable offspring when crossing with females (see below).

To confirm the XX karyotype of pseudomales, we crossed *Arh-mas-1* pseudomales (in APS4 background) with wildtype females (APS6 background). Using markers for the X chromosome, we genotyped pseudomales derived from self-progeny of F1 hybrid hermaphrodites. We found that *Arh-mas-1* pseudomales were heterozygous for the X chromosome markers (Fig. 2F, samples 1-3), confirming that this is a strain with an XX karyotype.

### *Arh-mas-1* XX pseudomales produce 0-4X sperm

In crosses between XX pseudomales and females, most of the F1 offspring is hermaphrodite or female (Fig. 3A), indicating that most of the *Arh-mas-1* spermatogenesis events generate viable 1X sperm. Cytological studies indicate that there is high variation in the way the X chromosomes segregate (Fig. 3B). In contrast to *C. elegans* (Fig. 1B), we never observed a lagging chromosome in anaphase I of *A. rhodensis* wildtype XO males. Instead, the unpaired X chromosome lags in anaphase II (Fig. 1B, Fig. 3B)(Shakes et al., 2011). Although the majority (66%, N=224) of the primary spermatocytes in XX pseudomales (*Arh-mas-1*) segregate DNA symmetrically, we also observed primary spermatocytes that segregate unequal amounts of DNA to the daughter cells (33%, N= 224). These occasional asymmetric cell divisions are supported by the observation of lagging X chromosomes during anaphase I (Fig. 3B). The different sizes of lagging chromosome masses could be due to (1) segregation of homologous X chromosomes to the same side, and (2) the premature separation of X chromatids in anaphase I (see models in Fig. 2A).

**Figure 3.**
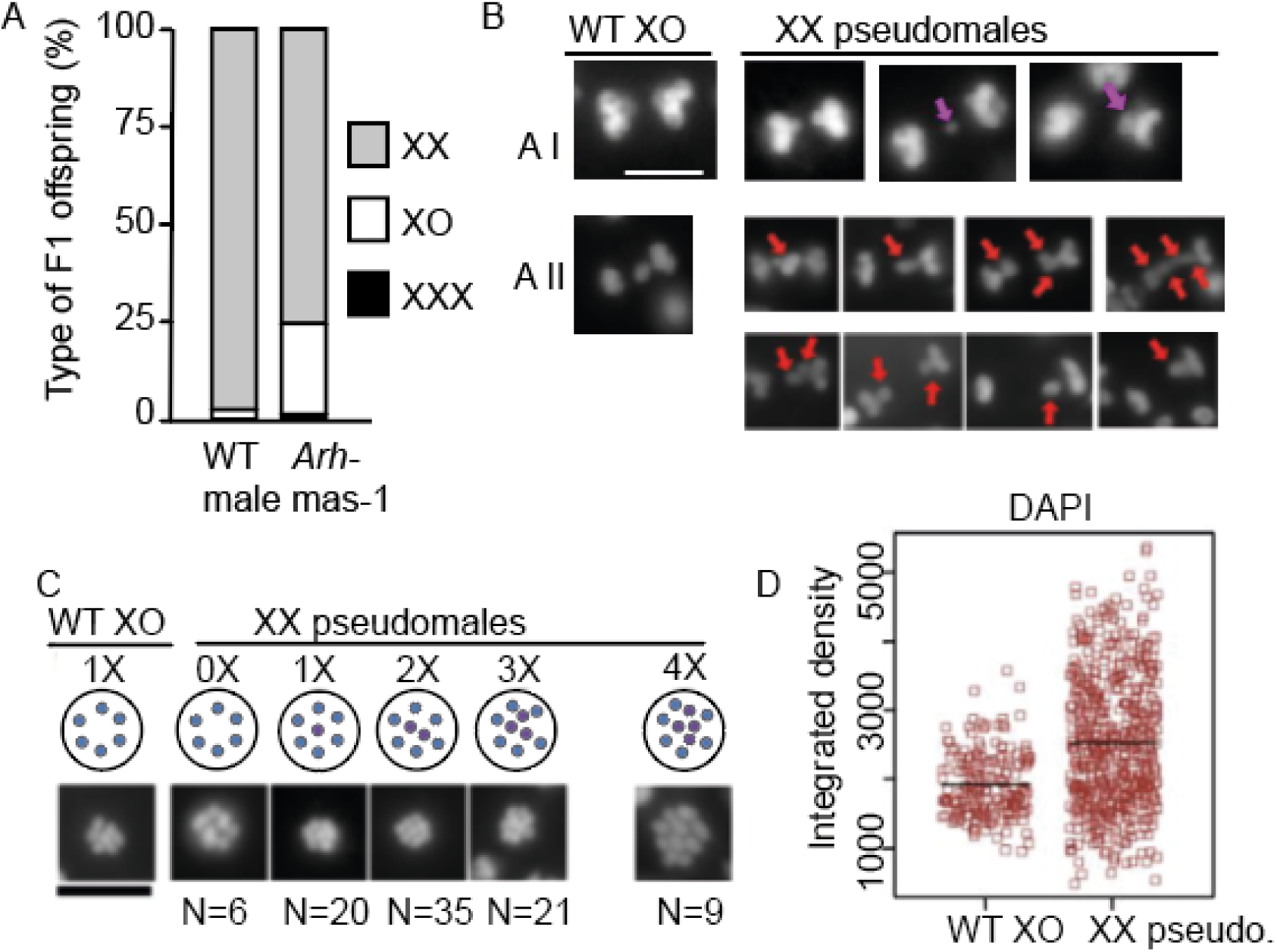
Types of F1 offspring after crossing *Arh-mas-1* pseudomales with females. **(A)** Bar graph representing the proportion of XX animals (females or hermaphrodites), XO (males) and XXX (dumpy). 19 crosses were performed between APS4 wildtype males and females (N= 5629 offspring) and 23 crosses were performed between *Arh-mas-1* pseudomales and APS4 females (N= 3135 offspring). (**B)** Meiosis in wildtype *A. rhodensis* XO males and XX pseudomales. The pink arrow is a lagging chromosome (stained with Hoechst) in anaphase I (A I) and red arrows indicate lagging chromosomes (stained with DAPI) in anaphase II (A II). Scale bar= 5 μm. (**C**) A variable number of chromosomes in metaphase plates, stained with DAPI. N= number of spermatocytes observed for each category, out of a total of 91. (**D**) Quantification of DAPI in MSP-stained sperm.

During anaphase II in spermatocytes of XX pseudomales, lagging chromosomes are also segregated symmetrically (59%, N= 54) (Fig. 3B) or asymmetrically (35%, N= 54)(Fig. 3B). Chromosome-lagging has been attributed to the attachment of microtubules to both sides of an unpaired chromosome, delaying its segregation to one of the poles (Fabig et al., 2020).

About 1% of the F1s from XX pseudomales and females were dumpy animals (Fig. 3A), possibly as the result of the fertilization of XX-sperm with an X-bearing oocyte. These dumpy animals are unlikely to be the result of background mutations because the strain was backcrossed three times and because we have not observed dumpy phenotype offspring from hermaphrodites harbouring the *Arh-mas-1* mutation. In *C. elegans*, XXX animals have a dumpy phenotype because of dosage compensation defects (Hodgkin, 1979; Vargas et al., 2017).

About a quarter of the offspring of a cross between a XX *Arh-mas-1* pseudomale and a XX female is XO male (Fig. 3A). Those F1 XO males are likely the result of the fertilization between an X-bearing sperm and a nullo-oocyte. The formation of a nullo-oocyte in females is rare, but it occurs in ∼3% of the meiosis (Shakes et al., 2011). Confirming this hypothesis, we found that 23% (N= 57) of the tested F1 males harbour the paternal X chromosome (Fig. 2F, samples 4, 5). An alternative scenario for the formation of XO offspring is the fertilization of an X-bearing oocyte by a nullo-X sperm. This seems to have been the case for the remaining 77% (N=57) of the cases (Fig. 2F, sample 6), in which F1 males inherited the maternal X chromosome.

To determine more directly the number of chromosomes in sperm, we examined spermatocytes stained with the DNA-staining dyes DAPI or Hoechst 33342. In metaphase plates, in which the number of chromosomes can be quantified, we confirmed the previous observation that *A. rhodensis* XO males have 6 autosomes and 1X chromosome (Fig. 3C) (Shakes et al., 2011). In XX pseudomales we observed cells only with autosomes (0X), or with X chromosomes that varied in number between 1-4 (Fig. 4C). To identify mature, functional sperm, we used the cytoskeleton protein Major Sperm Protein (MSP) as a marker. In those cells, the range of DNA intensity and size of those sperm was broader than in wild-type XO males (Fig. 3D), confirming the variability in the number of chromosomes in XX pseudomales. These variations reflect models 1-4 (Fig. 2A) and are useful to test if the X chromosome guides the partitioning of cytoplasmic components (see below).

**Figure 4.**
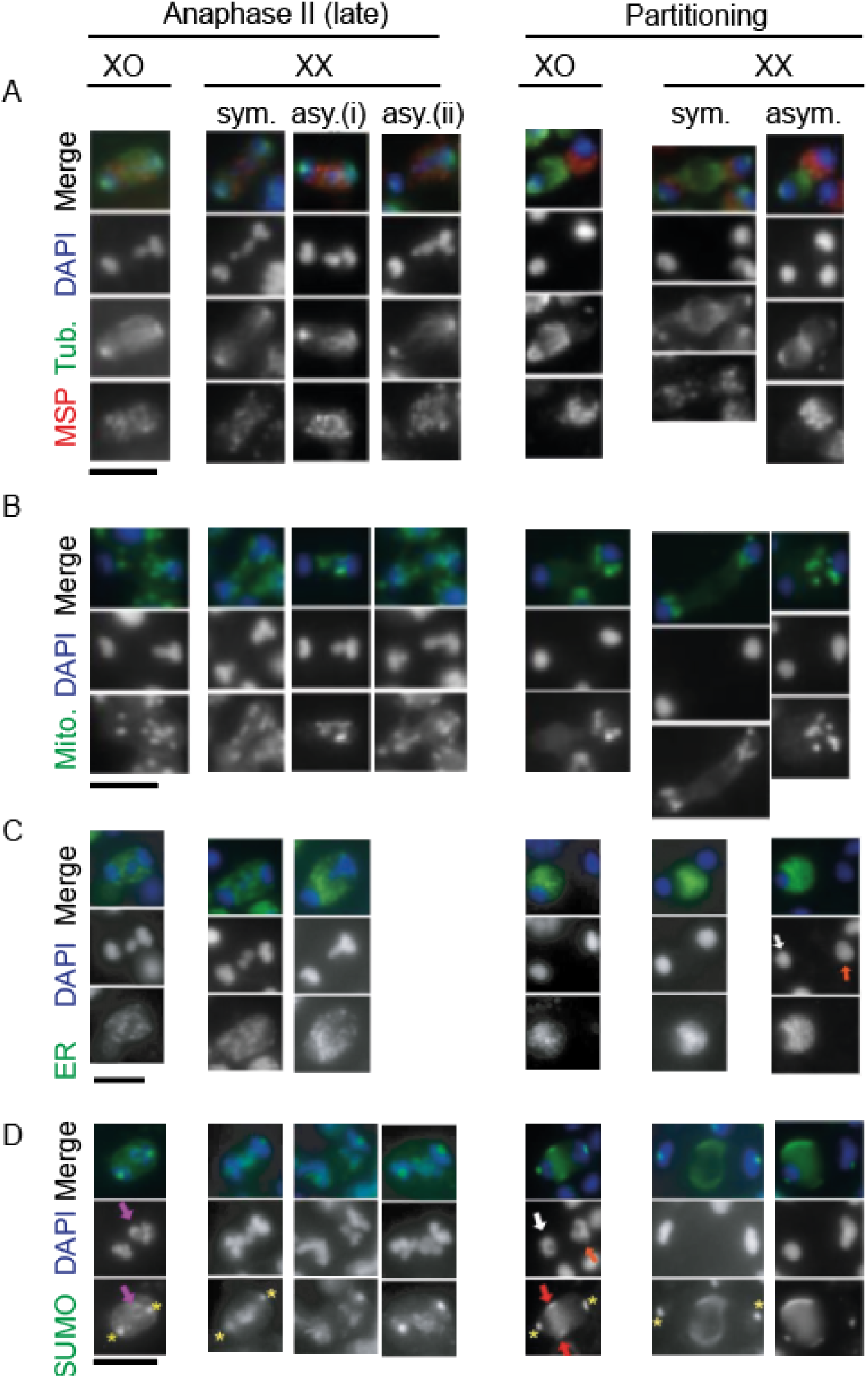
Asymmetric distribution of cytoplasmic components. Sperm spreads of XO wildtype males and XX pseudomales stained with antibodies against alpha-tubulin (**A**), MSP (**A**), mitochondrial beta-subunit of ATP synthase (**B**), ER-located cyp-33E1 (cytochrome P450 family) (**C**) and SUMO (**D**). In cells dividing asymmetrically (asym.), they may have one (i) or more chromosomes (ii) migrating to opposite poles. The orange and white arrows indicate larger and smaller DNA masses, respectively. The red arrows for the SUMO antibody staining indicate the expanding cortical bands. The yellow asterisks are centrosomes. Scale bar= 5 μm.

### Post-meiotic sperm components co-segregate with the X chromosomes

To examine the pattern of segregation of cytoplasmic components relative to the X chromosome segregation, we performed immunostainings on fixed sperm spreads. We chose cytoplasmic components that are essential for post-meiotic sperm (MSP and mitochondria) and others that are discarded into residual bodies (endoplasmic reticulum, alpha-tubulin and Small Ubiquitin-like Modifier (SUMO)). MSP is a protein contained within organelles called fibrous bodies in spermatocytes and is required in mature sperm for sperm motility (Smith, 2014). SUMO conjugation has been implicated in a variety of cellular processes (Drabikowski et al., 2018), including the regulation of meiotic proteins in *C. elegans* (Davis-Roca et al., 2018; Pelisch et al., 2017). Except for mitochondria and SUMO, we previously described the distribution of MSP, tubulin and endoplasmic reticulum (ER) in XO *A. rhodensis* spermatocytes (Shakes et al., 2011; Winter et al., 2017).

During the first meiotic division in wildtype XO males, there is symmetric segregation of X chromatids, autosomal homologous chromosomes and all five cytoplasmic components (Fig. S1A-D). In XX pseudomales, we observe the same pattern of bipolar distribution for all markers (Fig. S1A-D). Equal partitioning of cytoplasmic components occurs even when there is asymmetric segregation of chromosomes (63/194 primary spermatocytes), with one pole receiving more DNA than the other. During early anaphase II, when the lagging X is not yet in contact with autosomes, we also observe the equal distribution of cytoplasmic components in XO males and XX pseudomales for all markers (N= 407 secondary spermatocytes) (Fig. S1A-D).

In late anaphase II, when the X chromosome is biased towards one of the poles, asymmetries start to appear for most of the tested cytoplasmic components (Fig. 4). In wildtype XO animals, microtubules show a dense and elongated shape on the pole with the X chromosome (Fig. 4A). Similarly, mitochondria and MSP start to show a polarization towards the side with the X chromosome. The ER, however, tends to stay in a more central position of the dividing cell (Fig. 4C). SUMO antibody staining includes labelling of the lagging X chromosome, which is distinctly asymmetric (Fig. 4D).

The post-meiotic partitioning stage has the most pronounced asymmetries of cytoplasmic components. By then, all components are distributed in a unipolar pattern, with MSP and mitochondria at the pole with the highest DNA content, whereas tubulin, ER and SUMO with the opposite pattern (Fig. 4A-D). The pattern of distribution of SUMO in the cortex of the cell to become the residual body is like the previously described distribution of actin (Winter et al., 2017).

In XX pseudomales, X chromosomes segregate symmetrically in 30% of the cases (N=829), with both poles containing equal amounts of DNA. The remaining cells (70%, N=829), which segregate DNA asymmetrically, may have one or more chromosomes segregating to one of the poles. With no exceptions, we observe that MSP distribution is located around the centre of the symmetrically dividing cells in the late anaphase II and partitioning phase, but it is unipolar with a bias towards the side with more DNA in asymmetrically dividing cells (Fig. 4A). Similarly, the mitochondria distribute equally in cells that divide the chromosomes symmetrically, but with a bias towards the pole with more DNA (Fig. 4B). In instances in which there are lagging chromosomes on both sides, but with unequal amounts of DNA (Fig. 4B), mitochondria partition to both sides but shows a higher prevalence in the pole with more DNA.

For the components that are discarded into polar bodies in wildtype XO males (tubulin, ER, SUMO), in XX pseudomales they follow the predicted patterns of distributing equally or centrally in cells that divide with equal amounts of DNA. In asymmetric segregation of DNA, these components partition to the pole with less DNA. In some of the cells with symmetric amounts of DNA segregation, tubulin is located centrally in the partitioning phase (Fig. 4A). This is reminiscent of residual body formation in *C. elegans* (Ward et al., 1983), which forms centrally in this organism but never in wildtype *A. rhodensis* (Shakes et al., 2011). The ER distribution in XX pseudomales remains at the centre of symmetrically dividing cells in both anaphase II and partitioning phase (Fig. 4C). In cells with asymmetric segregation of chromosomes, the ER distribution concentrates in the pole with less DNA (Fig. 4C). The SUMO distribution in late anaphase II reflects in large part the staining of lagging chromosomes, which can be equally partitioned in cells dividing with equal amounts of DNA or biased towards one of the poles in asymmetrically dividing cells (Fig. 4D). When partitioning, SUMO localization is in the central part of the cortex of symmetrically dividing cells, or in the cortex of the cell with less DNA (Fig. 4D).

## DISCUSSION

The initial observation of a few male offspring derived from *Auanema* male and female crosses is the result of a modification in spermatogenesis: males discard male-generating spermatids (nullo-X sperm) and promote the production of X-bearing sperm (Shakes et al., 2011; Winter et al., 2017). This is a mechanism that generates biased sex ratios, which provides adaptive advantages in certain ecological circumstances (Van Goor et al., 2021). Here we provide support for the hypothesis that the X chromosome acts as a signal for an ACD to generate viable X-bearing sperm cells. The X chromosome would be an example of an intrinsic signal for polarising cell divisions.

Asymmetric segregation of proteins, RNAs and organelles during cell division occurs in prokaryotes and eukaryotes (Horvitz and Herskowitz, 1992; Inaba and Yamashita, 2012; Morrison and Kimble, 2006; Sunchu and Cabernard, 2020). In mitotic cells, biased distribution of cytoplasmic components may result in cells with different cell fate determinants, organelles and protein aggregates. Previously reported examples for intrinsic cues include random events (Broadus and Doe, 1997) and co-segregation with organelles that are intrinsically asymmetric (Chen and Yamashita, 2021).

This type of cell division found in *Auanema* spermatogenesis, in which residual material is partitioned to a cell that contains chromosomes, is atypical in metazoans (Gorelick et al., 2016). Residual bodies, which are often observed in ACDs during spermiogenesis, are usually devoid of DNA (Gorelick et al., 2016; Zakrzewski et al., 2021). There are a few exceptions, such as in the nematode *Rhabdias*, in which one of the X chromosomes is discarded into a residual body during the hermaphrodite spermatogenesis (Runey et al., 1978). During residual body formation In *C. elegans* spermatogenesis, motor proteins (SPE-15, myosin VI; NMY-2, non-muscle myosin II)(Hu et al., 2019; Kelleher et al., 2000), and a cargo adaptor protein (GIPC, RGS-GAIP-interacting protein C) (Hu et al., 2019) have been implicated in the correct segregation of organelles. However, the polarisation signal for this asymmetric segregation is unknown.

Contrary to the canonical meiosis, the unpaired X chromosome in *Auanema* males separates its sister chromatids already in the first meiotic division (Shakes et al., 2011; Winter et al., 2017). The result is that each secondary spermatocyte has an X chromosome. Thus, during cell division of these cells, there is potential for the X chromosome to act as a polarising signal. Such a scenario does not occur in *C. elegans* males (Fig. 1), as this nematode follows the canonical meiosis (reviewed in (Chu and Shakes, 2013)): each secondary spermatocyte will either have the X chromosome or not. When the secondary spermatocytes divide, each daughter cell will contain the same chromosomal content (either with an X or without) as the parental cell. Therefore, the polarisation mechanisms for the correct partition of cytoplasmic content for the residual bodies is likely to be different in *C. elegans*.

The mechanism by which the X chromosome may act as a polarising cue in *Auanema* is still unclear. It is possible, for instance, that a secondary nucleation centre for cytoskeleton proteins, such as non-centrosomal microtubule-organising centres, emanate from the X chromosome (Sanchez and Feldman, 2017). Further genetics studies, as well as electron microscopy and live-cell imaging will help to elucidate the possible mechanisms.

## ACKNOWLEDGEMENTS

A.P.-d.S. was supported by a grant from Leverhulme Trust (RPG-2019-329). The *E. coli* strain was provided by the CGC, which is funded by NIH Office of Research Infrastructure Programs (P40 OD010440). D.C.S. was supported by a grant from the National Science Foundation (<u>IOS 1122101</u>).

---

**Supplemental Figure 1.**
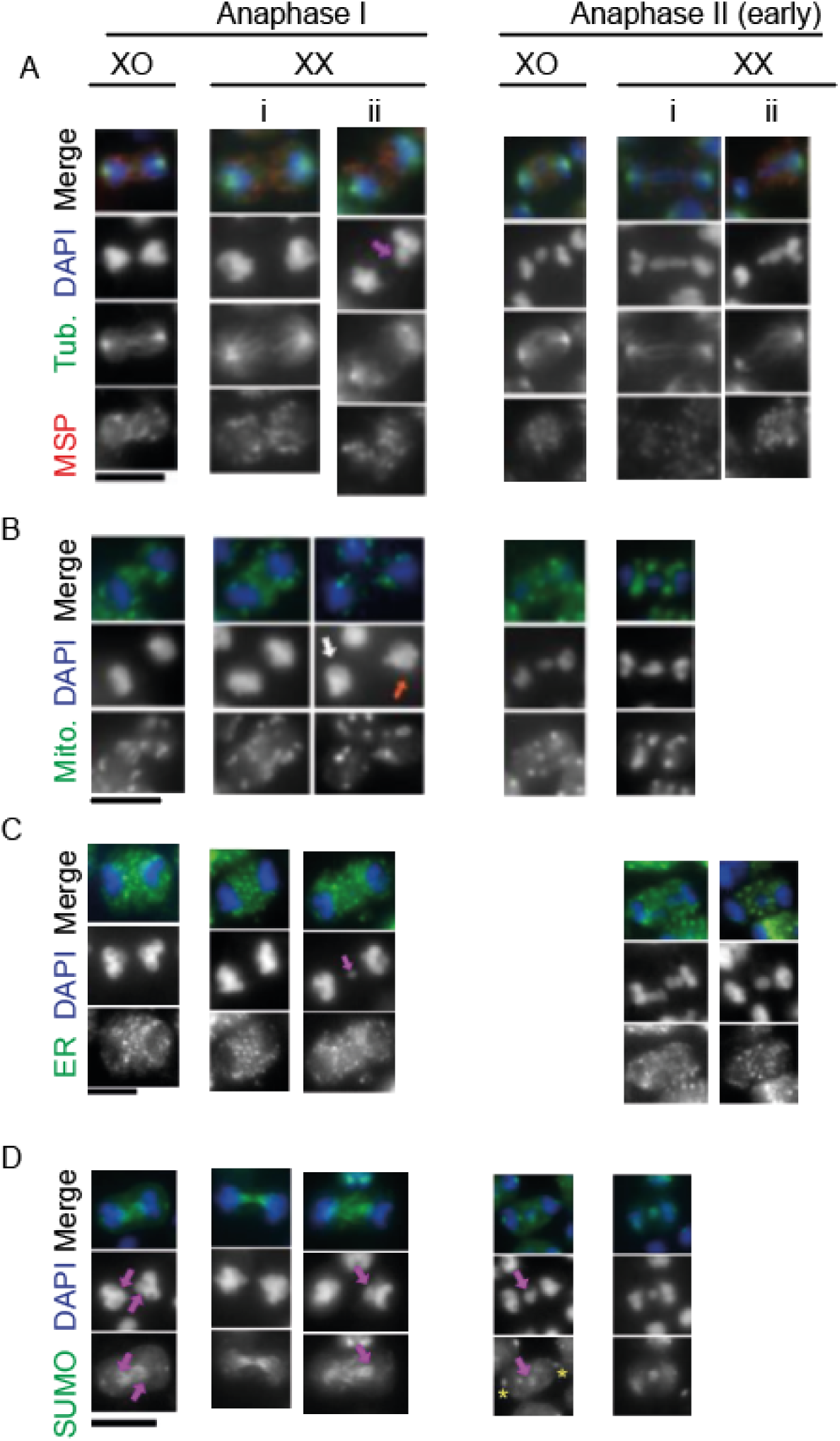
Cytoplasmic components segregate symmetrically in anaphase I and early anaphase II. In wildtype XO males and XX pseudomales, the cytoplasmic components MSP (A), alpha-tubulin (A), mitochondria (B), endoplasmic reticulum (ER) (C) and SUMO (D) distribute equally to both poles of the dividing spermatocytes. The pink arrow indicates lagging chromosomes, whereas the red and white arrows indicate larger and smaller DNA masses, respectively. Scale bar= 5 μm.

## Notes

### Competing Interest Statement

The authors have declared no competing interest.

